# Alternative electron pathways of photosynthesis drive the algal CO_2_ concentrating mechanism

**DOI:** 10.1101/2021.02.25.432959

**Authors:** Adrien Burlacot, Ousmane Dao, Pascaline Auroy, Stephan Cuiné, Yonghua Li-Beisson, Gilles Peltier

**Author notes:** correspondence to Gilles Peltier. Howard Hughes Medical Institute, Department of Plant and Microbial Biology, 111 Koshland Hall, University of California, Berkeley, CA 94720-3102 USA. **Author contributions**: A.B. and G.P. designed the research; A.B., O.D., P.A., S.C., and G.P. performed research; A.B. and G.P. contributed new reagents/analytic tools; A.B. and G.P. analyzed data; A.B. and G.P. wrote the paper with inputs from Y.L.-B.

## Abstract

Global photosynthesis consumes ten times more CO_2_ than net anthropogenic emissions, and microalgae account for nearly half of this consumption^1^. The great efficiency of algal photosynthesis relies on a mechanism concentrating CO_2_ (CCM) at the catalytic site of the carboxylating enzyme RuBisCO, thus enhancing CO_2_ fixation^2^. While many cellular components involved in the transport and sequestration of inorganic carbon (C_i_) have been uncovered^3,4^, the way microalgae supply energy to concentrate CO_2_ against a thermodynamic gradient remains elusive^4-6^. Here, by monitoring dissolved CO_2_ consumption, unidirectional O_2_ exchange and the chlorophyll fluorescence parameter NPQ in the green alga *Chlamydomonas*, we show that the complementary effects of cyclic electron flow and O_2_ photoreduction, respectively mediated by PGRL1 and flavodiiron proteins, generate the proton motive force (*pmf*) required by C_i_ transport across thylakoid membranes. We demonstrate that the trans-thylakoid *pmf* is used by bestrophin-like C_i_ transporters and further establish that a chloroplast-to-mitochondria electron flow contributes to energize non-thylakoid C_i_ transporters, most likely by supplying ATP. We propose an integrated view of the CCM energy supply network, describing how algal cells distribute photosynthesis energy to power different C_i_ transporters, thus paving the way to the transfer of a functional algal CCM in plants towards improving crop productivity.

**One sentence summary:** Photosynthetic alternative electron flows and mitochondrial respiration drive the algal CO_2_ concentrating mechanism

## Introduction

In aquatic ecosystems, microalgal photosynthesis has to cope with a low CO_2_ availability resulting from the slow diffusion of CO_2_ in water^7^. Moreover, the CO_2_-fixing enzyme Ribulose-1,5-Bisphosphate Carboxylase-Oxygenase (RuBisCO) has a low affinity for CO_2_^8^, thus the efficiency of algal photosynthesis strongly relies on a CO_2_-Concentrating Mechanism (CCM)^9^. The algal CCM involves the sequential actions of inorganic carbon (C_i_) transporters and carbonic anhydrases located in different cellular compartments^10^, and results in active accumulation of CO_2_ at the RuBisCO level^4,6^. Several CCM components have been identified in the green alga *Chlamydomonas reinhardtii* (*Chlamydomonas* hereafter)^3,4^, particularly putative C_i_ transporters operating across the plasma membrane (High Light Activated 3, HLA3)^11^, the chloroplast envelope (Low Carbon Inducible A, LCIA)^12,13^, and more recently across the thylakoid membrane (bestrophin-like transporters, BSTs)^14^. The transport of C_i_ across membrane bilayers against a concentration gradient is an energy-dependent process^6,14^, and the role of photosynthesis in supplying the chemical energy required by the functioning of CCM has been early recognized^15^.

During photosynthesis, sunlight is converted into chemical energy by two photosystems (PSII and PSI) acting in series through the so-called Linear Electron Flow (LEF), reducing NADP^+^ into NADPH and producing a *pmf* across the thylakoid membrane. The *pmf* is then used for ATP synthesis, and both NADPH and ATP supply energy for CO_2_ fixation. However, LEF produces less ATP than required for CO_2_ fixation as compared to NADPH^16^, and photosynthesis relies on additional mechanisms to fill this gap^17^. These include *i*. cyclic electron flow around PSI (CEF), which involves both Proton Gradient Regulation-5 (PGR5)^18,19^ and Proton Gradient Regulation Like-1 (PGRL1) proteins in plants and algae^20,21^ and *ii*. pseudo-cyclic electron flow (PCEF) which diverts electrons to O_2_ at the PSI acceptor side^22^, catalyzed by flavodiiron proteins (FLVs) in cyanobacteria^23^, bryophytes^24,25^ and green microalgae^26^. Both CEF and PCEF generate a *pmf* without producing NADPH, thus re-equilibrating the high NADPH/ATP ratio of LEF. Another pathway involving several metabolic shuttles between chloroplast and mitochondrial respiration, called here chloroplast-to-mitochondria electron flow (CMEF), can also supply extra ATP for CO_2_ fixation when CEF is absent or deficient^27,28^. In this context, how photosynthesis energy is delivered to the different C_i_ transporters and how can this be done without compromising CO_2_ fixation capacity are pivotal questions^4,6^.

In this work, we addressed these questions by studying *Chlamydomonas* mutants of CEF (*pgrl1*^21^), PCEF (*flvB*^26^) and of the recently discovered thylakoid C_i_ transporters BSTs^14^. We show that CCM activity is unaffected in the single CEF or PCEF mutants but severely impaired in the double mutants and that the trans-thylakoid *pmf* produced by the cooperative action of these mechanisms is used by BST thylakoid C_i_ transporters. We further establish that CMEF is involved in CCM functioning, most likely by supplying ATP to plasma membrane and/or chloroplast envelope C_i_ transporters, thus revealing how transport steps distant from the thylakoid can be empowered.

## Results

### Combined deletion of FLVs and PGRL1 impairs C_i_ affinity and growth in air

To investigate the involvement of FLV-mediated PCEF and PGRL1-mediated CEF in the energy supply to the CCM, we first measured net photosynthetic O_2_ production at various C_i_ concentrations in *Chlamydomonas flvB*^26^ or *pgrl1*^21^ single mutants. When cells were grown under air (400 ppm CO_2_, low CO_2_), C_i_ affinities were similar for control WT strains and single mutants (K_1/2_ ∼10 µM), indicating a fully functional CCM (**Fig. 1 A, B, H**). Under high CO_2_ (3% CO_2_ supplementation to air), *flvB* and *pgrl1* mutants as well as their respective progenitors showed similar affinity for C_i_ with a K_1/2_ around 100 µM (**Fig. 1 D, E, H**). In order to assess possible functional redundancy between FLVs and PGRL1, double mutants were obtained by genetic crosses of the single *pgrl1* and *flvB* mutants (**Extended data Fig. 1**). Among the progenies, five independent double mutants were isolated (*pgrl1 flvB*-1, 2, 3, 4 and 5) as well as four independent control strains exhibiting normal accumulation of both FLVs and PGRL1 (thereafter called WT-1, 2, 3 and 4) (**Fig. 1 G; Extended data Fig. 1**). While no difference in C_i_ affinities was observed between these strains when grown at high CO_2_, double mutants showed a 7 times lower affinity for C_i_ as compared to control strains when grown at low CO_2_ (**Fig. 1 C, F, H; Extended data Fig. 2 B, D**).

**Figure 1.**
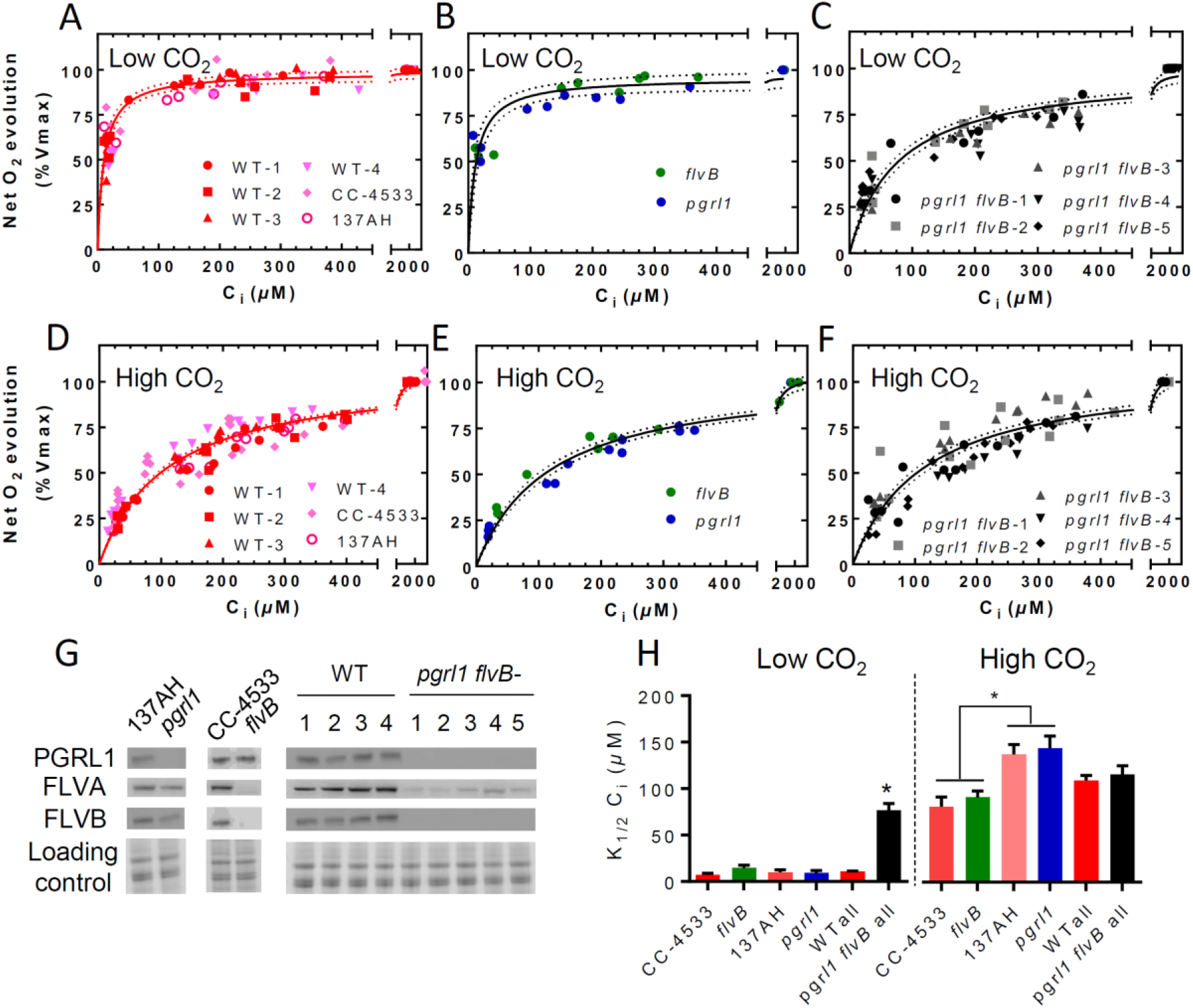
Affinity of photosynthetic O_2_ evolution for inorganic carbon (C_i_) is decreased in *pgrl1 flvB double* mutants, but unaffected in *flvB* and *pgrl1* single mutants. Net O_2_ production was measured at pH 7.2 in cells grown under 400 ppm CO_2_ air (Low CO_2_) (**A, B and C**) or 3% CO_2_ (High CO_2_) (**D, E and F**). Shown are three replicates for each strain (dots) and hyperbolic fit with variability (plain lines, dotted lines). For each replicate, net O_2_ production was measured following stepwise C_i_ addition, and normalized to the maximum photosynthetic net O_2_ production. Since these strains were generated in different genetic backgrounds (CC-4533 and 137AH, respectively) showing contrasted photosynthetic activities (**Extended data Fig. 2 A, C**), data shown are normalized to the maximal net O_2_ production rate. (**A, D**) 137AH and CC-4533 are the respective control strains for *pgrl1* and *flvB*, WT-1 to -4 are four independent control strains obtained from the *pgrl1* × *flvB* crossing. (**B, E**) *flvB* and *pgrl1* mutant strains. (**C, F**) *pgrl1 flvB*-1 to -5 are five independent double mutant strains. **G** Immunodetection of PGRL1, FLVA and FLVB in the different strains with Coomassie blue staining as the loading control. (**H)** K_1/2_ values as determined from the hyperbolic fit for each strain. Shown are mean ± SD (n=3 for single mutants and their controls), values for all double mutant strains have been pooled (“*pgrl1 flvB* all”, n=15) as well as their control strains (“WT all”, n=12). Asterisks represent significant differences (p<0.05, one way ANOVA with Tukey correction).

Mutants defective in the CCM often cannot grow properly in low CO_2_. Here, we compared growth at different CO_2_ concentrations, pH and light intensities. While all strains showed similar growth at high CO_2_, the growth of *pgrl1 flvB* double mutants was impaired under low CO_2_ and very low CO_2_ (100 ppm CO_2_ in air) (**Fig. 2 A**), similar to the growth defect observed in the CCM-deficient mutant *cia5* (**Fig. 2 A; Extended data Fig. 3**). The growth defect observed in double mutants worsened with light intensity but was barely affected by pH (**Extended data Fig. 3**). The accumulation of the major CCM components, as evaluated by immunodetection, was similar in all strains, with the exception of LCI1 which is present in lower amount in double mutants (**Fig. 2 B**) and to a lesser extent in single mutants (**Extended data Fig.4 C**). Since growth of the LCI1 knock-out mutant was shown to be not affected by low CO_2_ ^29^, we conclude however that the *pgrl1 flvB* mutants growth defect is not due to the lower LCI1 level. Carbonic anhydrase activity measured *in vivo* was induced in low CO_2_ reaching similar levels in all strains (**Fig. 2 C**). Double mutants and control strains showed similar maximal O_2_ photosynthetic production (**Extended data Fig. 2 A, C**) and PSII quantum yields (**Extended data Fig. 4 A,**), as well as similar levels of major photosynthetic complexes (**Extended data Fig. 4 B**). We conclude from these experiments that PGRL1-mediated CEF and FLV-mediated PCEF contribute to the CCM operation likely by supplying energy, and can compensate each other, as exemplified by the absence of phenotype in single mutants.

**Figure 2.**
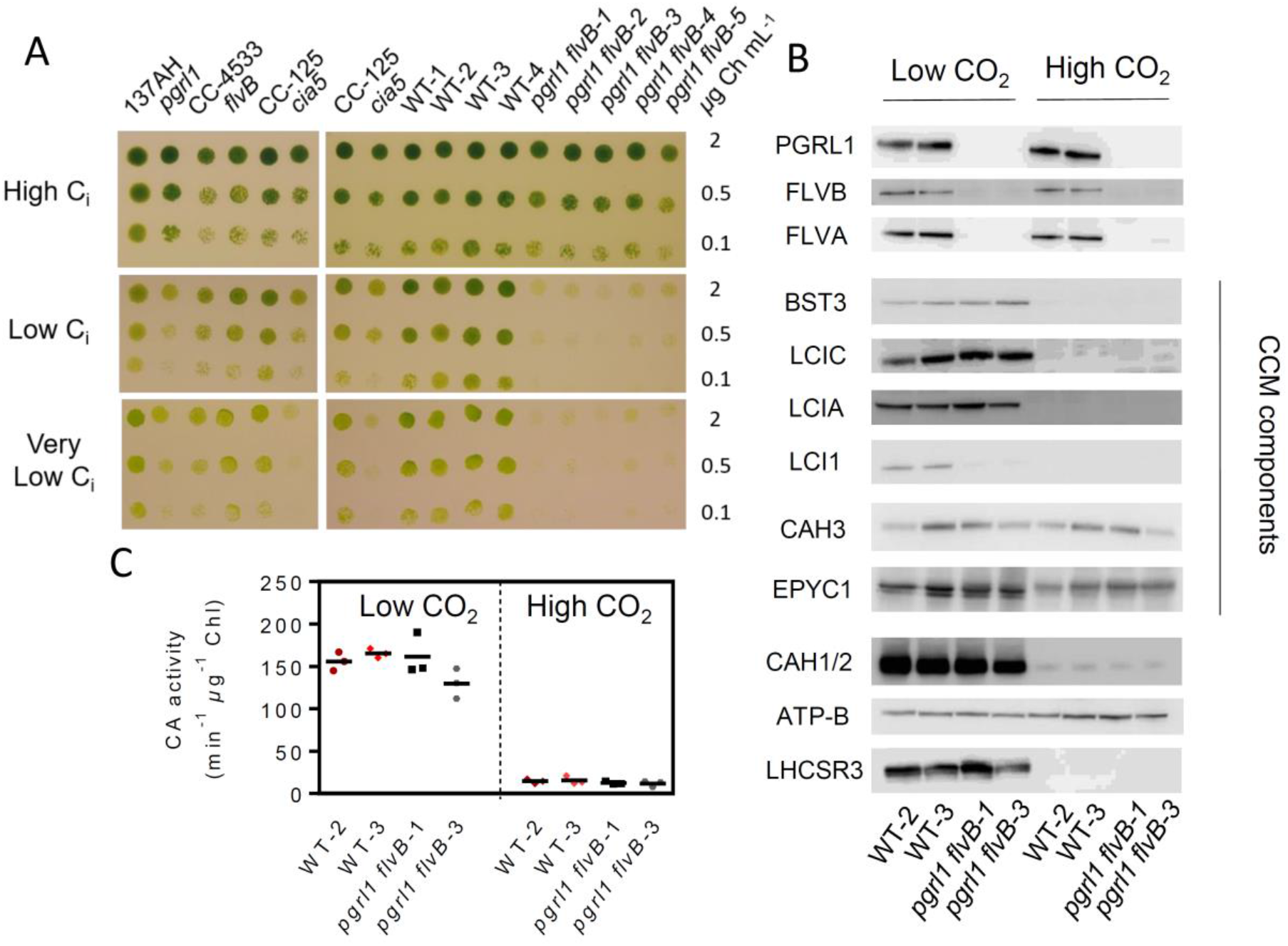
Growth of *pgrl1 flvB* double mutants is impaired at low CO_2_ while CCM components are present. **(A)**. Growth tests performed on *pgrl1, flvB*, and their corresponding control strains (137AH and CC-4533 respectively) (left panels) and on double mutants (*pgrl1 flvB*-1 to -5) and their control strains (WT-1 to -4) (right panels); the CCM1 mutant *cia5* was introduced as a CCM-deficient control together with its reference strain CC-125. Cells were spotted on plates containing minimal medium at pH 7.2 and grown under continuous illumination (60 µmol photon m^-2^ s^-1^) either under High CO_2_, Low CO_2_ or Very Low CO_2_ (100 ppm CO_2_ in air). Shown are representative spots of ten independent experiments. **(B)** Immunodetection of PGRL1, FLVA, FLVB and of the major CCM components in two independent *pgrl1 flvB* double mutants and controls grown in Low CO_2_ or High CO_2_. (**C)** Carbonic anhydrase (CA) activity was determined *in vivo* by following the unlabelling of ^18^O-enriched CO_2_ in the same strains and conditions as in **B**. Shown are mean values and replicates (n=3).

### The *pmf* generated by CEF and PCEF energizes the CCM at the level of C**_i_**transport across the thylakoid membrane mediated by BSTs

To gain further insight into the link between CEF, PCEF and CCM energization, we assessed the level of the *pmf* dynamics across the thylakoid membranes in the different mutant strains during the functioning of the CCM. Currently, no direct measurement of *pmf* is available, but it can be inferred by the level of the rapidly reversible non-photochemical-quenching component (qE). The Light-Harvesting Complex Stress-Related 3 protein (LHCSR3), responsible for qE is activated by low luminal pH, making qE a sensitive and reliable probe of luminal pH^30,31^ (**Fig. 3 A**). We thus took advantage of the presence of LHCSR3 in low CO_2_ -grown cells (**Fig. 2 B, Extended data Fig. 4 C**) to quantify the luminal pH and the *pmf*. In both control lines and single mutants, the NPQ level was maximal when C_i_ level was low, and rapidly and reversibly decreased either in the light upon C_i_ injection or at low C_i_ when light was turned off (**Fig. 3 B; Extended data Fig. 5 A-E, I**). In sharp contrast, no CO_2_ -dependent NPQ could be observed in the *pgrl1 flvB* mutants (**Fig. 3 C; Extended data Fig. 5 F, J**) where only a slowly inducible and irreversible NPQ was observed. We conclude from this experiment that CEF and PCEF energize the CCM through the generation of a trans-thylakoidal *pmf*, both mechanisms being able to substitute for each other.

**Figure 3.**
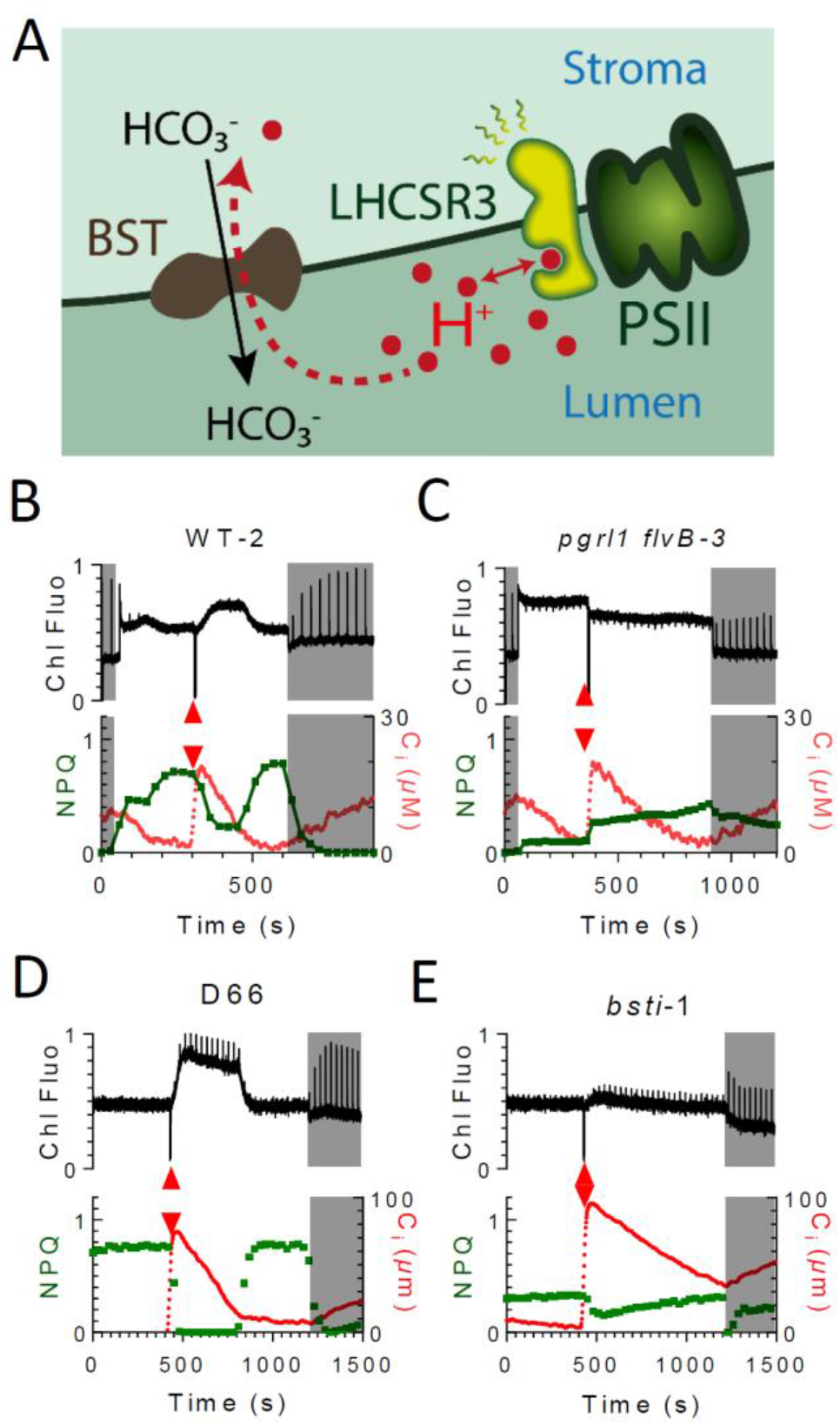
The NPQ dependence to C_i_ concentration is abolished in *pgrl1 flvB* double mutants and in the BST mutant *bsti*-1. (**A)** Schematic view describing the rationale of the experiment; NPQ, which depends on the luminal pH (via LHCSR3) is used to probe the trans-thylakoidal *pmf* during the CCM functioning. (**B-E**) Combined measurements of chlorophyll fluorescence (upper panels), C_i_ concentrations and NPQ (lower panels) during dark-light-dark transients in WT-2 (**B**), *pgrl1 flvB-3* (**C**), the *bsti*-1 control strain D66 (**D**) and *bsti*-1 (**E**). All strains were grown at low CO_2_. Shown are representative experiments (n=3). Red arrows indicate addition of bicarbonate.

Recently, three BST-like proteins were proposed to transport C_i_ at the thylakoid level^14^. To gain insight into the role of the *pmf* in the supply of energy to BSTs, we assessed the kinetics of the C_i_-dependent NPQ in a BST knock-down strain (*bsti*-1)^14^. While the control strain showed a C_i_-dependent NPQ (**Fig. 3 D)**, the NPQ of *bsti*-1 was barely affected during C_i_ depletion (**Fig. 3 E; Extended data Fig. 5 G, H)**, thus indicating that the *pmf* is not consumed during C_i_ depletion in the *bsti*-1 mutant. The low NPQ levels in *bsti-*1 are attributable here to the lower accumulation of LHCSR3 as compared to its control strain (**Extended data Fig. 4 D,**). We conclude from this experiment that the *pmf* generated by CEF and PCEF is used by BST proteins to transport C_i_ across the thylakoid membranes (**Fig. 3 A**).

### CMEF is the third protagonist of the triumvirate

In order to gain quantitative insight into the nature of compensation mechanisms in *pgrl1*, we investigated the C_i_ dependence of the light-dependent O_2_ consumption measured using ^18^O-labelled O_2_. As previously reported^32,33^, O_2_ consumption increased at low C_i_ in control strains (**Extended data Fig. 6 A, E)**. In *pgrl1*, O_2_ uptake rates were higher than in control strains (**Extended data Fig. 6 A-D**), which was previously attributed to an increase of the FLV-mediated PCEF^27^. O_2_ uptake rates were strongly diminished in *flvB*^26^, but surprisingly, a C_i_-dependent O_2_ uptake process remained (**Extended data Fig. 6 E, F**). In order to determine whether mitochondrial respiration is responsible for the remaining light-dependent O_2_ uptake, we used two mitochondrial respiration inhibitors, myxothiazol and salicyl hydroxamic acid (SHAM), which inhibit the cytochrome *bc*_*1*_ complex and the mitochondrial alternative oxidase, respectively. We show that the remaining light-dependent O_2_ uptake measured at low C_i_ in *flvB* is indeed due to mitochondrial respiratory activity driven by photosynthesis and therefore attributed to CMEF (**Extended data Fig. 6 K**).

We then investigated the contribution of mitochondria to CCM energization in wild-type strains. While addition of respiratory inhibitors had no effect on the V_Max_ and C_i_ affinity of high CO_2_ -grown strains (K_1/2_∼100 µM) (**Fig. 4 A, Extended data Fig. 7 D-F, I**), it reduced by half the C_i_ affinity of low CO_2_ -grown control strains (K_1/2_>20 µM) as compared to untreated cells (K_1/2_ ∼10 µM) (**Fig. 4 B**). The effect on C_i_ affinity was observed only when myxothiazol and SHAM were added together (**Extended data Fig. 8**), indicating that both alternative oxidase and cytochrome *bc*_*1*_ electron pathways contribute to CCM energization. Importantly, respiratory inhibitors also increased the K_1/2_ of air-grown *bsti*-1 mutant (**Fig. 4 B, Extended data Fig. 7 H**), thus showing that the contribution of mitochondria to CCM energization operates at the level of other transporters than thylakoidal BSTs. The contribution of the different pathways (CEF, PCEF and CMEF) was deduced from O_2_ consumption rates measured in the different mutants during C_i_ depletion (**Extended data Fig. 6**). While the contribution of CEF remained relatively constant, the contribution of FLV-mediated PCEF and of CMEF dramatically increased at low C_i_ (**Fig. 5 A)**.

**Figure 4.**
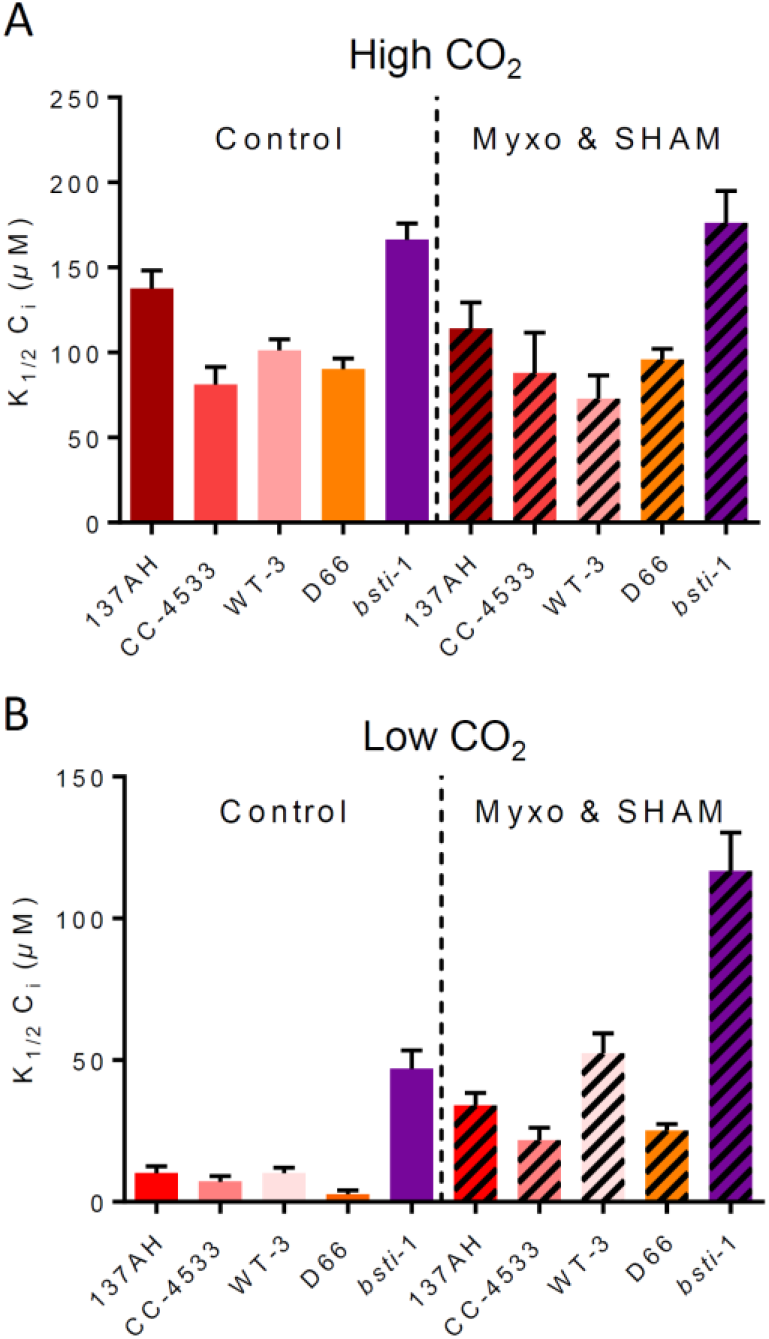
Mitochondrial inhibitors decrease the affinity of photosynthesis for C_i_ in low CO_2_ -grown cells. Photosynthetic net O_2_ production was measured as in **Fig. 1** in High CO_2_ (**A**) or Low CO_2_ (**B**) grown cells. K_1/2_ values were determined from hyperbolic fits in the different strains in the presence or absence of two mitochondrial inhibitors myxothiazol (Myxo, 2.5 *µ*M) and salicyl hydroxamic acid (SHAM, 400 *µ*M), respectively acting on the cytochrome *bc*_*1*_ complex and on the alternative oxidase. Shown are mean values (n=3, ±SD). All strains showed K_1/2_ significantly different when treated with myxothiazol and SHAM as compared to their control when grown in low CO_2_ (p<0.05; one way ANOVA with Tukey correction).

## Discussion

Although the CCM’s requirement for photosynthesis energy has been early recognized^15^, the associated molecular mechanisms have remained poorly understood^4^. The participation of PCEF^33^ or CEF^34^ has been proposed, but their actual and respective contributions have not been established. In this work, we demonstrate that both FLV-mediated PCEF and PGRL1-mediated CEF cooperate to supply energy to the CCM. Moreover, we show that the trans-thylakoidal *pmf* generated by both mechanisms is the energy vector used by the recently discovered trans-thylakoidal BST-like C_i_ transporters^14^.

We further establish that mitochondrial respiration, the role of which in CCM energization has so far been largely ignored, provides energy to other CCM transporters than BSTs, through an efficient inter-organelles redox trafficking. From the analysis of the respective contribution of each mechanism as a function of C_i_ concentration, we conclude that while CEF contribution is relatively constant, PCEF contribution increases at low C_i_ concentrations, and that contribution of CMEF also becomes important at the lowest C_i_ concentrations. Since these are typical conditions where putative ATP-dependent periplasmic and chloroplast envelope transporters (HLA3 and LCIA respectively) are highly active^4,12,13^, we propose that the ATP produced by CMEF at low C_i_ supplies energy to one or both transporters (**Fig. 5**). Interestingly, LCI1 accumulation, whose function is tightly linked to HLA3^3,29^, decreased in *pgrl1, flvB, pgrl1 flvB* and *bsti-1*, indicating that impairments of C_i_ transport at the thylakoid level may regulate periplasmic transport processes mediated by LCI1. This could be due to an increased cytosolic C_i_ concentration resulting from the absence of functional thylakoid C_i_ transport, which would in turn trigger down-regulation of LCI1 expression to avoid cytosolic C_i_ over-accumulation.

**Figure 5.**
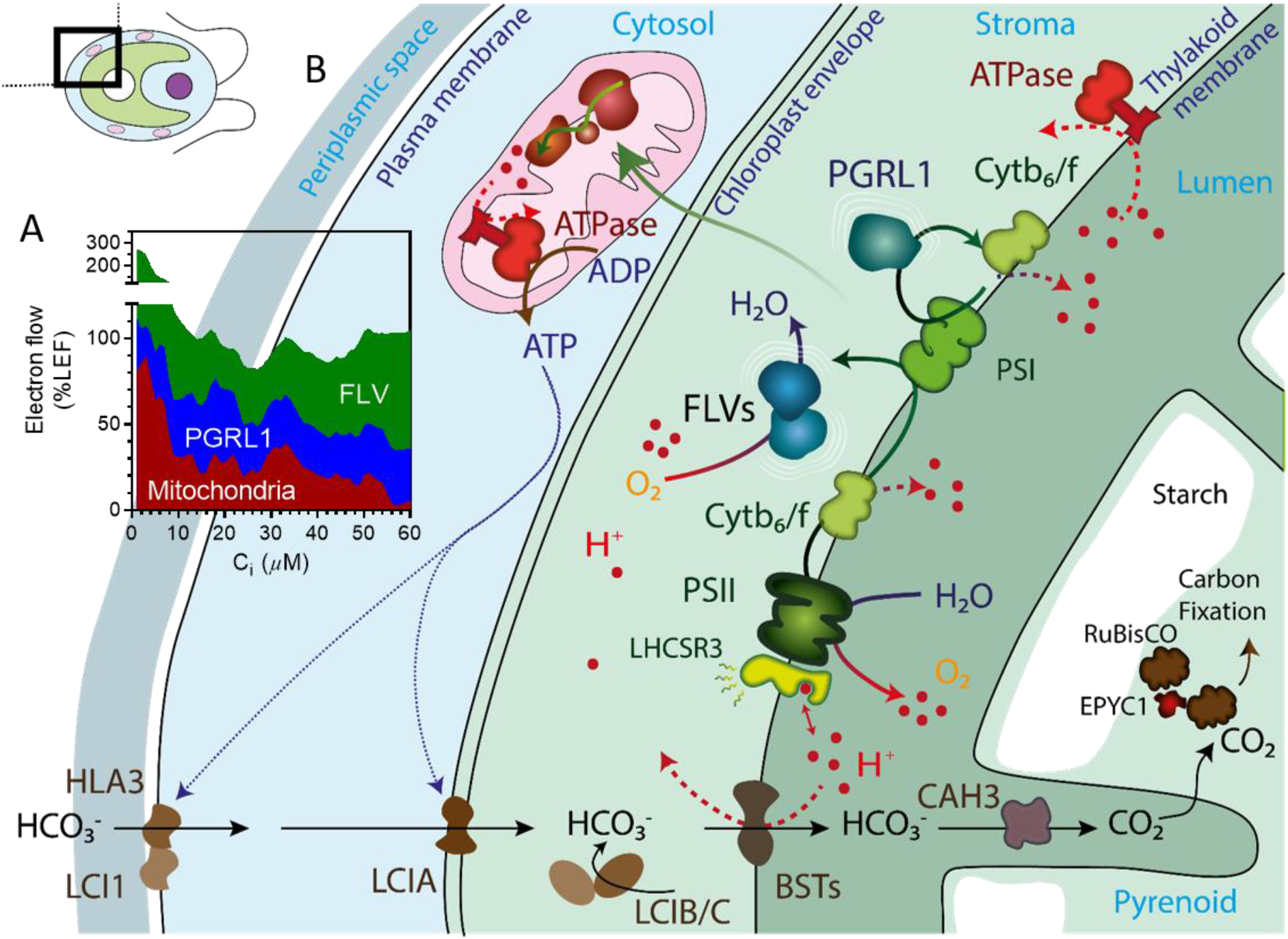
Proposed mechanism of CCM energization network in algal cells. (A) The relative contribution of FLV-mediated PCEF, PGRL1-mediated CEF and CMEF to CCM energization at different C_i_ concentrations are quantified from O_2_ exchange measurements performed in the different mutant strains and expressed as a percentage of LEF (**Extended data Fig. 6**). (**B)** Schematic view of the energy supply network to the CCM. CCM components including LCI1, HLA3, LCIA and BSTs transporters, LCIB, LCIC, CAH3, EYPC1 and RuBisCO are shown in brown, and components of the photosynthetic electron transport chain (PSII, PSI, Cytb_6_/f) in green. We propose here that energization of BSTs-dependent C_i_ transport is mediated by the *pmf* produced by the combined action of FLVs-mediated O_2_ photoreduction and the PGRL1-mediated cyclic electron flow. CMEF would generate the ATP needed to power other cellular C_i_ transporters such as LCIA and HLA3.

The presence of an active CCM is a key factor influencing phytoplankton biomass production in the oceans^2^, especially for phytoplankton species producing large oceanic blooms^35^. However, the lack of knowledge on CCM operation *in situ*^9^ makes it difficult to predict how global changes will affect phytoplankton communities^36^. We demonstrate here that the presence of a functional CCM can be probed by measuring C_i_-dependent NPQ, which could be used as a simple parameter to determine CCM activity in aquatic ecosystems. In diatoms which are important in marine ecosystems^37^ and well-studied species^38^, FLVs are absent (they are restricted to the green algal lineage), and CEF is working at very low levels^28^, and it was proposed that the ATP requirement of photosynthesis was fulfilled by an efficient chloroplast to mitochondria energy trafficking^28^. Based on the dependence of CCM on mitochondrial respiration observed here, we propose that in diatoms such energy trafficking between chloroplast and mitochondria could play an important role in fulfilling the ATP requirement of CCM transporters, including BST-like transporters which are well conserved in most microalgal phyla^39^.

A potential limitation of a thylakoid C_i_ pump consuming the *pmf* would be competition with the synthesis of ATP required for CO_2_ fixation^14^. This is particularly critical since LEF is known to supply less ATP than required for CO_2_ fixation^17^. We suggest here that the combined action of the three alternative mechanisms, CEF, PCEF and CMEF, which all result in an increase of the ATP/NADPH ratio, allows to fulfill the energy requirement of the CCM without compromising CO_2_ fixation. A major biotechnological challenge in CCM research is the improvement of crop productivity by transferring microalgal components into higher plants^40-42^. Building a fully functional CCM in plants represents a tremendous scientific challenge, which has recently regained considerable interest^4,43^. Our study shows that an integrated understanding of the cellular energetics is key towards fulfilling the energy requirement of a synthetic CCM without compromising the efficiency of photosynthetic CO_2_ fixation. For instance, the expression of FLVs in higher plants which has been shown to supply extra *pmf* during photosynthesis^44-47^, appears as a promising starting point to supply the extra energy needed to power thylakoid C_i_ transporters of the BST family in plants. We foresee that future research coupling energy source and CCM expression should help boost plant productivity.

## Accession numbers

Genes studied in this article can be found on https://phytozome.jgi.doe.gov/ under the loci Cre12.g531900 (FLVA), Cre16.g691800 (FLVB), Cre07.g340200 (PGRL1), Cre16.g662600 (BST1), Cre16.g663400 (BST2), and Cre16.g663450 (BST3).

## List of abbreviations

BST: Bestrophin-like proteins
CCM: Carbon Concentrating Mechanism
CEF: Cyclic Electron Flow
C_i_: Inorganic Carbon
CMEF: Chloroplast-Mitochondria Electron Flow
HLA3: High Light Activated 3
LCI: Low Carbon Induced
LEF: Linear Electron Flow
LHCSR3: Light Harvesting Complex Stress Related 3
Myxo: Myxothiazol
NPQ: Non Photochemical Quenching
PCEF: Pseudo-Cyclic Electron Flow
PGRL1: Proton Gradient Regulation Like-1
FLVs: Flavodiiron proteins
PSI: Photosystem I
PSII: Photosystem II
RuBisCO: Ribulose Bisphosphate Carboxylase Oxygenase
SHAM: Salicyl hydroxamic acid.

## Materials and Methods

*Chlamydomonas flvB, pgrl1, bsti*-1 mutants and their respective wild-type CC-4533, 137AH and D66 were previously described^14,21,26^. All strains were grown phototrophically under moderate light (80 µmol photons m^-2^ s^-1^) in minimal medium either under low CO_2_ or high CO_2_. Gas exchange rates were measured using a membrane inlet mass spectrometer^48^ and combined NPQ measurements were done as previously described^49^. All replicates shown are biological replicates from independent cultures. Other methods are described in Supplementary Materials and Methods.

## Acknowledgments

This work was supported by the A*MIDEX (ANR-11-IDEX-0001-02) project and by the ANR OTOLHYD. Ousmane Dao is the recipient of a PhD grant awarded to Y. L-B. The authors thank Pr. Graham Peers for stimulating discussion, Pr. Krishna K. Niyogi and Dr. Masakazu Iwai for critical reading of the manuscript, Pr. James V. Moroney for providing the *bsti-1* mutant, Pr. Luke Mackinder and Pr. Hideya Fukuzawa for respectively providing BST3, and LCIA, LCI1 antibodies. Contributions of Dr. Solène Moulin for artistic drawings of Fig. 5, of Stephanie Blangy for LCIC antibody preparation, Emma Calikanzaros for preliminary experiments and Arthur Gosset for performing genetic crosses of *flvB* and *pgrl1* mutant, are gratefully acknowledged. The authors acknowledge the European Union Regional Developing Fund, the Region Provence Alpes Côte d’Azur, the French Ministry of Research, and the CEA for funding the HelioBiotec platform.

## Data and materials availability

All data needed to evaluate the conclusions in the paper are present in the paper and/or the Supplementary Materials.

