## Supplemental Material and Methods for "Alternative electron pathways of photosynthesis drive the algal CO_2_ concentrating mechanism"

### Supplemental Materials and Methods for: Alternative electron pathways of photosynthesis drive the algal $CO_2$ concentrating mechanism

Adrien Burlacot*, Ousmane Dao, Pascaline Auroy, Stephan Cuiné, Yonghua Li-Beisson, Gilles Peltier+

Aix Marseille Univ, CEA, CNRS, Institut de Biosciences et Biotechnologies Aix-Marseille, CEA Cadarache, 13108 Saint-Paul-lez-Durance, France

*present address: Howard Hughes Medical Institute, Department of Plant and Microbial Biology, 111 Koshland Hall, University of California, Berkeley, CA 94720-3102 USA

**ORCID IDs:** 0000-0001-7434-6416 (A.B.), 0000-0002-7040-5770 (O.D.); 0000-0002-3376-6550 (P.A.), 0000-0002-3000-3355 (S.C.), 0000-0003-1064-1816 (Y.L.-B.), 0000-0002-2226-3931 (G.P.)

***Chlamydomonas* strains and cultivation.** All *Chlamydomonas* strains were grown photo-autotrophically in 150 mL flasks at 25°C in a buffered minimal medium (20 mM MOPS pH= 7.2) under constant illumination (80 µmol photons $m^{-2}$ $s^{-1}$). Cells were harvested during the exponential phase. The $mt^{+}$ *flvB-21* mutant generated in [1] was crossed with the $mt^{-}$ *pgrl1* mutant, and the progenies were screened based on chlorophyll fluorescence transients for *FLVB* insertion and PCR for *PGRL1* insertion (**Extended data Fig. 2 B and C**). Five independent strains exhibiting a *flvB* mutant-like chlorophyll fluorescence transient [2] and harboring an insertion of the paromomycin cassette at the *PGRL1* locus were selected (**Extended data Fig. 2 D**). Four independent strains exhibiting wild-type chlorophyll fluorescence phenotype and no insertion at the *PGRL1* locus were selected as control strains (**Extended data Fig. 2 D**). The absence of FLVB and PGRL1 proteins in the five independent double mutants (named *pgrl1 flvB-1, -2, -3, -4,* and *-5*) and their presence in the four independent control strains (named WT-1, -2, -3 and -4) was confirmed by immunodetection (**Fig. 1 G**). The *bsti*-1 mutant and its control strain D66 was described in [3].

**DNA extractions and PCR amplification.** Total DNA was extracted by mixing one cell colony in 50 µL of a 10 mM Na-EDTA (Sigma-Aldrich) solution. Putative insertion in the *PGRL1* locus was confirmed in progeny of the crossing by PCR using *dreemTaq* DNA polymerase with green GC Buffer (Thermo Scientific). The two primer sequences used to amplify the *PRGL1* locus were: 5'-TTACGCAGCGGCCTTAGCCTTCTTCTTGGC-3' and 5'-GGCTAAGCGCTGTGTGCCGC-3'. PCR cycles were: 2 min at 95°C / 40 cycles: 30 s at 95°C, 30 s at 60°C, 2 min at 72°C / 4 min at 72°C for final extension. PCR products were separated on 1% (w/v) agarose gels.

**Immunoblot analysis.** For protein analysis, $10^6$ of exponentially growing cells (≈10 µg chlorophyll $mL^{-1}$) were harvested by centrifugation at 4,000 g for 3 min at 4°C. Pellets were resuspended in 200 µL of 1% SDS, 800 µL of cold acetone were added in order to extract chlorophyll and incubated for 30 min at -20°C. Samples were then centrifuged for 10 min, 16,000 *g* at 4°C and chlorophyll concentration measured on the supernatant. The protein pellet was resuspended with Novex™ Nupage™ LDS buffer 1x (Invitrogen™), and proteins were then denaturated at 70°C for 20 min. Protein extracts (10 µg protein) were loaded on Novex™ Nupage™ Bis tris 12% (Invitrogen™) gel, run 1 h at 190 V in Novex™ Nupage™ (Invitrogen™) MOPS buffer or MES (for PsaC) buffers and transferred to nitrocellulose membrane (or PVDF for LCIC) using semidry transfer technique. Immunodetection was performed using antibodies raised against PGRL1 [4], FLVB and FLVA [5], NDA2 [6]. Other antibodies against PsbD (AS06 146), PsaC (AS10 939), Cytb6 (AS03-034), RbcL (AS03 037), ATP-B (AS05-085), COXIIB (AS06 151), AOX1 (AS06 152), FeSOD (AS06-125), EPYC1 (AS09-602), LHCSR3 (AS14-2766), CAH1/ 2 (AS11-1737) and CAH3 (AS05 073) were obtained from Agrisera (https://www.agrisera.com/). The LCIC antibody was generated against a synthetic peptide containing the sequence of 12 amino acids found in the C-terminus of LCIC as described in [7]. Two rabbits were injected with the synthetic peptide for the production of LCIC antibody. BST3 antibody is a kind gift from Pr. Luke Mackinder (University of York, England), LCI1 and LCIA antibodies are a kind gift of Pr. Hideya Fukuzawa (Kyoto University, Japan).

**Growth tests.** The different *Chlamydomonas* strains were cultivated at 400 ppm $CO_2$ (Low $CO_2$), except *cia5* and CC125 which were grown under 3% $CO_2$ (High $CO_2$) and acclimated to low $CO_2$ 24 hours before spotting as described in [8] because the *cia5* mutant is unable to grow autotrophically without High $CO_2$ [9]. Cells were harvested during exponential growth and resuspended in fresh minimal medium to 0.1, 0.5, or 2 µg chlorophyll $mL^{-1}$. Eight-microliter drops were spotted on plates at pH=7.2 or 8.2 (buffered with 20 mM MOPS or 20 µM Tris respectively) and exposed to High $CO_2$, Low $CO_2$ or Very Low $CO_2$. Homogeneous light was supplied by a panel of fluorescent tubes, and neutral filters were used to obtain the desired light intensity. Temperature was maintained at 25°C at the level of plates by means of fans.

**$CO_2$ affinity of net $O_2$ photosynthesis.** $CO_2$ affinity was determined using membrane inlet mass spectrometry (MIMS) [10]. Cells grown at air level or 3% $CO_2$ were harvested, centrifuged at 450 *g* for 3 min and resuspended in fresh minimal medium (pH=7.2) at 10 mg chlorophyll $mL^{-1}$. The cell suspension was then bubbled in the MIMS reaction vessel with a $CO_2$ depleted gas mixture (80 % $N_2$, 20 % $O_2$). Upon $CO_2$ depletion, the reaction vessel was closed, light was turned on (2,000 µmol photon $m^{-2}$ $s^{-1}$, green LEDs) and gas exchange recorded. Increasing amounts of bicarbonate were then sequentially added to reach various $C_i$ concentrations inside the reaction vessel during gas exchange measurements.

**Carbonic anhydrase activity measurements.** Carbonic anhydrase (CA) activity was determined in intact cells by monitoring $^{18}O/^{16}O$ isotope exchange between $^{18}O$-enriched $CO_2$ and $H_2O$ using MIMS as described in [10]. Doubly labelled $^{13}C^{18}O_2$ was prepared by equilibrating 1 mol $L^{-1}$ $NaH^{13}CO_3$ (99 % $^{13}C$-atom, Euriso-top, France) with $H_2{}^{18}O$ (97 % $^{18}O$-atom, Cambridge Isotope Lab. Inc.) for 24 h at room temperature. The reaction vessel (1.5 mL) contained minimal growth medium buffered with 20 mM MOPS (pH 7.2). At t = 0, 15 µL of a 1 mol $L^{-1}$ $NaH^{13}C^{18}O_3$ solution were added and the isotope exchange was measured by continuously recording the concentration of $^{13}C^{18}O^{18}O$ (m/z = 49), $^{13}C^{18}O^{16}O$ (m/z = 47), $^{13}C^{16}O^{16}O$ (m/z = 45). $CO_2$ unlabelling was followed by determining the time constant of the isotope content decrease [10-12], first for 3 min in the absence of algae, and then for 3 min following addition of the algal sample (20 µL at 50 µg chlorophyll $mL^{-1}$). Total carbonic anhydrase activity of the algal sample was expressed on a chlorophyll content basis, as the time constant of the isotope content decrease, after subtracting the exchange activity measured in the absence of algae.

**Chlorophyll fluorescence and NPQ measurements.** Chlorophyll fluorescence was measured using a Pulsed Amplitude Modulation (PAM) fluorimeter (Dual-PAM 100, Walz GmbH, Effeltrich, Germany) on the MIMS chamber as described in [2] using green actinic light (2,000 µmol photon $m^{-2}$ $s^{-1}$, green LEDs). Red saturating flashes (8,000 µmol photons $m^{-2}$ $s^{-1}$, 600 ms) were delivered to measure $F_M$ (in dark-acclimated samples) and then every 30 s to measure $F_M'$ (upon actinic light exposure). NPQ was calculated as $(F_M - F_M')/F_M'$.

**Calculations of electron fluxes though alternative pathways depending on $C_i$ concentration.** $O_2$ exchange was measured by MIMS using $^{18}O$-enriched $O_2$ as described in [10] on *pgrl1*, *flvB* and their respective control strains 137AH and CC-4533, strains. Upon illumination, the $C_i$ was depleted by photosynthesis, allowing to determine gas exchange rates at various $C_i$ concentrations [13]. $O_2$ exchange rates were averaged from three biological replicates (**Extended Data Fig. 6**). Since the CCM functioning in not affected in *pgrl1* (**Fig. 1**), we considered that the increase in $O_2$ uptake rates between *pgrl1* and its control strain ($O_2$ difference$_{pgrl1}$) (**Extended Data Fig. 6**) reflects the additional electron flux to $O_2$ compensating the absence of PGRL1-mediated CEF. As CEF is 1.5 × less efficient than PCEF to generate a proton gradient [14], CEF was calculated as equal to 1.5 × $O_2$ difference$_{pgrl1}$. The FLV-mediated PCEF was calculated as the difference in $O_2$ uptake rates between CC-4533 and *flvB* (**Extended Data Fig. 6**). CMEF was calculated as the light-induced $O_2$ uptake remaining in the *flvB* mutant (**Extended Data Fig. 6**). In order to determine the contribution of alternative electron pathways to CCM energizing as compared to $CO_2$ fixation, CEF was normalized to net $O_2$ evolution measured in *pgrl1* and both PCEF and CMEF were normalized to net $O_2$ evolution in *flvB*.
