## Supplemental figures for "Alternative electron pathways of photosynthesis drive the algal CO_2_ concentrating mechanism"

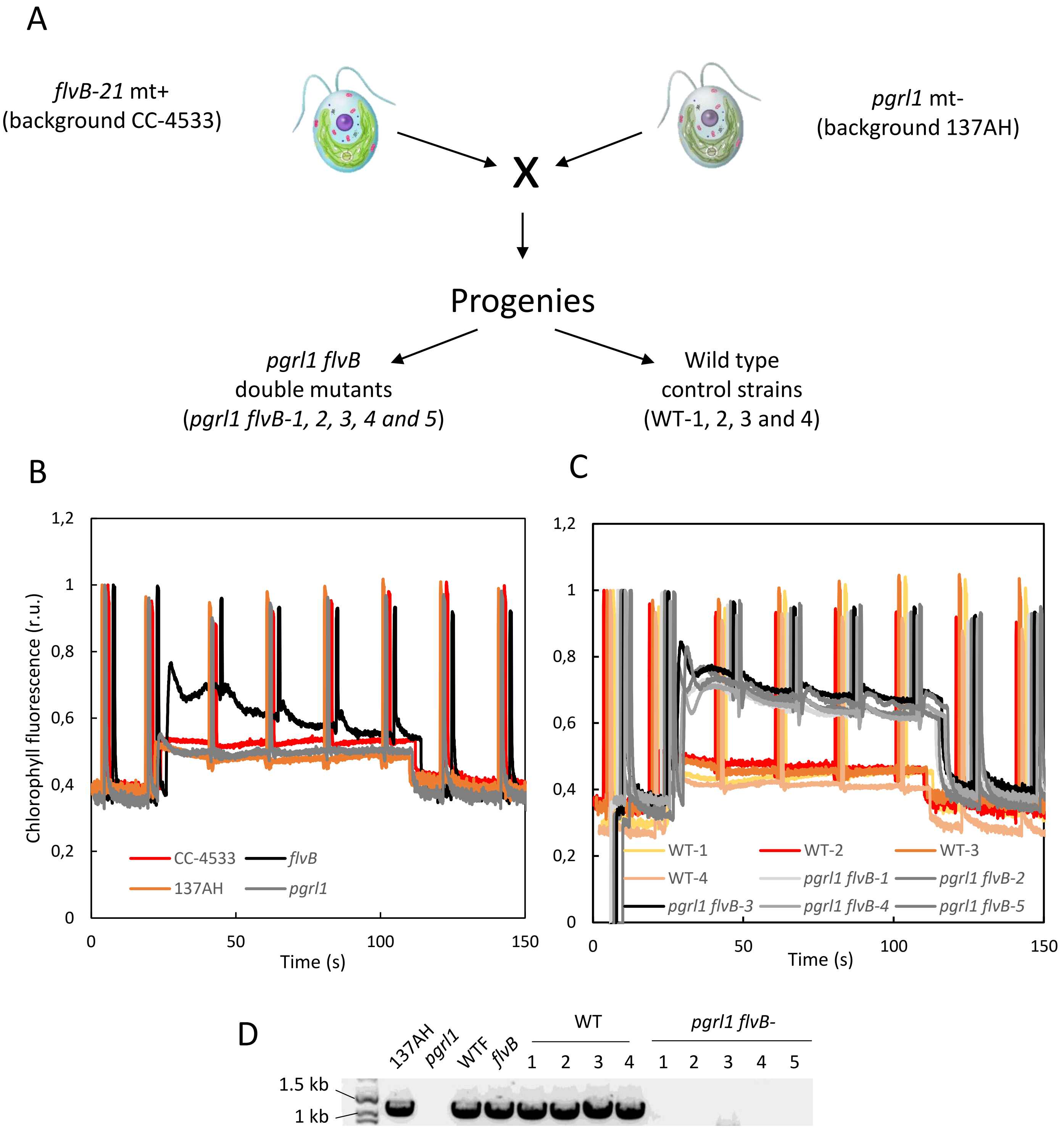


**Extended data Fig. 1. Obtaining and characterizing of *pgrl1 flvB* double mutants.** Upon crossing of *pgrl1* and *flvB*-21 opposite mating types, progenies were screened for FLVB deficiency based on the chlorophyll fluorescence pattern, and for insertion in the *PGRL1* locus by PCR. Five independent double mutants and control strains (with normal chlorophyll fluorescence pattern and no insertion at the *PRGL1* locus) were randomly selected (**A**). (**B, C)** Chlorophyll fluorescence patterns of parental strains (*pgrl1* and *flvB*-21) and of their respective control strains (137AH and CC-4533 respectively) in **B**, and of progenies of the crossing (*pgrl1 flvB*-1, 2, 3, 4 and 5; WT-1, 2, 3, 4) in **C**. Chlorophyll fluorescence measurements were performed using a PAM fluorimeter in the dark and under red actinic light (100 µmol photon $m^{-2}$ $s^{-1}$, from t=22 s to t=110 s). Data are normalized to initial $F_M$ and slightly shifted on the time axis for clarity. (**D)** PCR amplification targeting the *PGRL1* locus in the different strains.

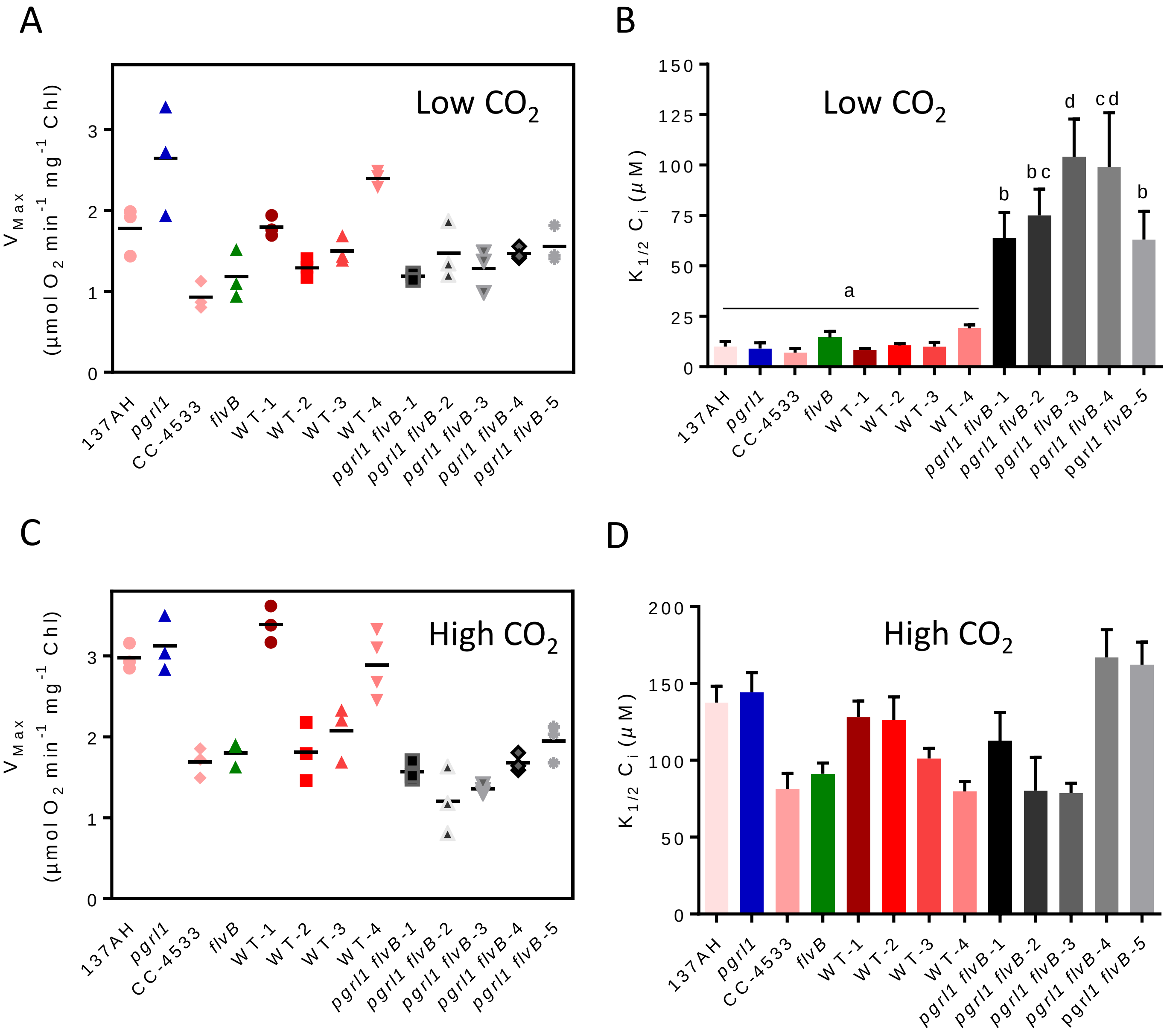


**Extended data Fig. 2. Maximum net $O_2$ evolution and $K_{1/2}$ values measured in *pgrl1*, *flvB*, *pgrl1 flvB* and their respective controls when grown under Low $CO_2$ or High $CO_2$.** Net $O_2$ production rates were measured as described in **Fig. 1** in low $CO_2$ (**A, B**) or high $CO_2$ (**D, E**) grown cells. (**A, C**) Maximal net $O_2$ production rates. Shown are mean values and replicates (n=3). (**B, D**) $K_{1/2}$ values determined for each strain from hyperbolic fits. Shown are mean ± SD (n=3). Letters (a, b, c, d) above bars in (**B**) represent significant differences ($p<0.05$) between strains based on ANOVA analysis.

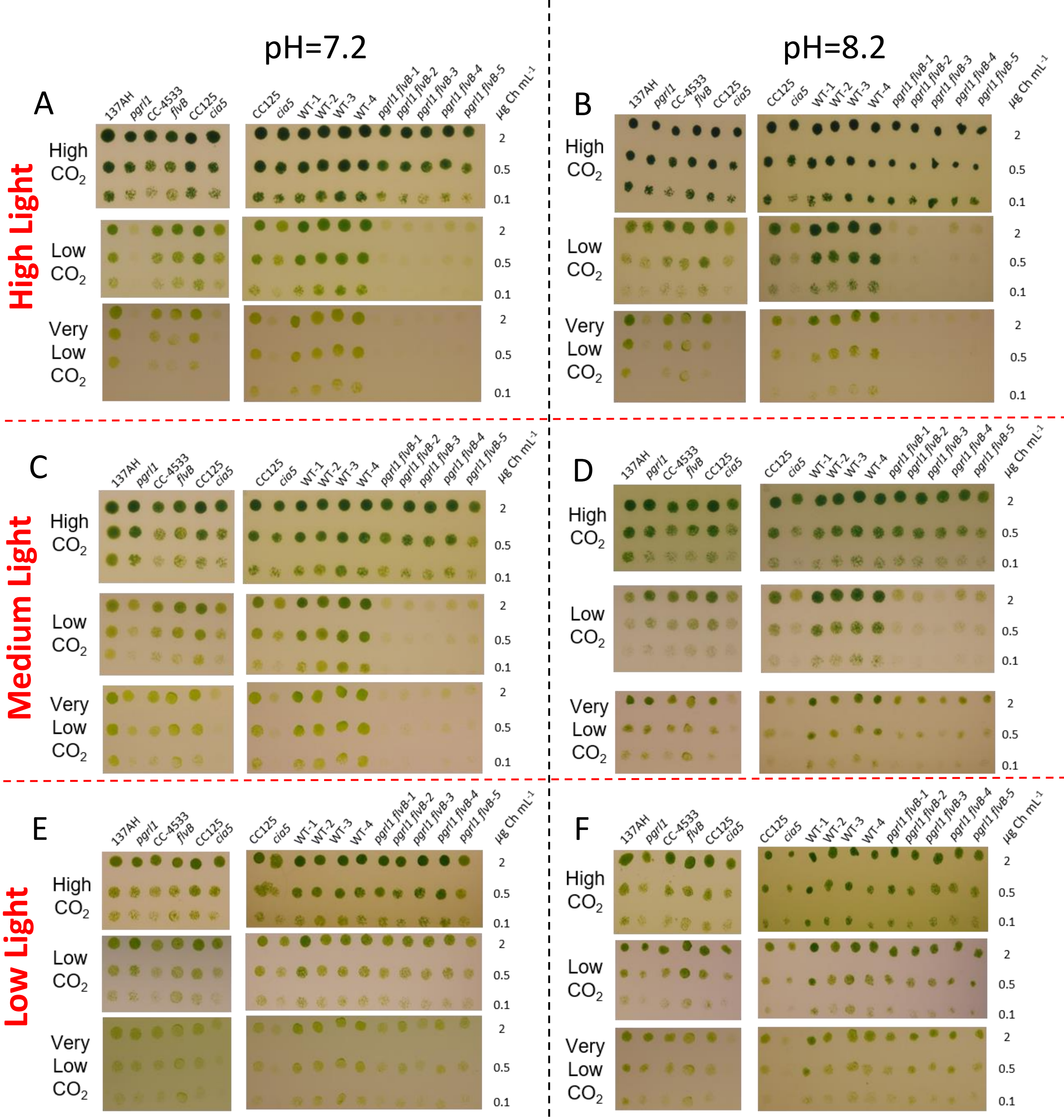


**Extended data Fig. 3. Growth of *pgrl1*, *flvB*, *pgrl1 flvB* mutants and their control strains.** Cells were spotted on plates containing minimal medium at pH 7.2 (**A, C, E**) or pH 8.2 (**B, D, F**) and grown under continuous low light (30 µmol photon $m^{-2}$ $s^{-1}$, **E, F**), medium light (60 µmol photon $m^{-2}$ $s^{-1}$, **C, D**) or high light (100 µmol photon $m^{-2}$ $s^{-1}$, **A, B**). For each pH and light conditions, cells were grown either at 100 ppm $CO_2$ (Very low $CO_2$), 400 ppm $CO_2$ (Low $CO_2$) or 3% of $CO_2$ (High $CO_2$). Growth was assessed in *pgrl1*, *flvB*, and their respective control strains (137AH and CC-4533) (left panels) and on double mutants (*pgrl1 flvB*-1 to -5) and their control strains (WT-1 to -4) (right panels); the CCM1 mutant *cia5* was introduced as a CCM-deficient control together with its reference strain CC-125. Shown are representative spot tests of ten independent experiments.

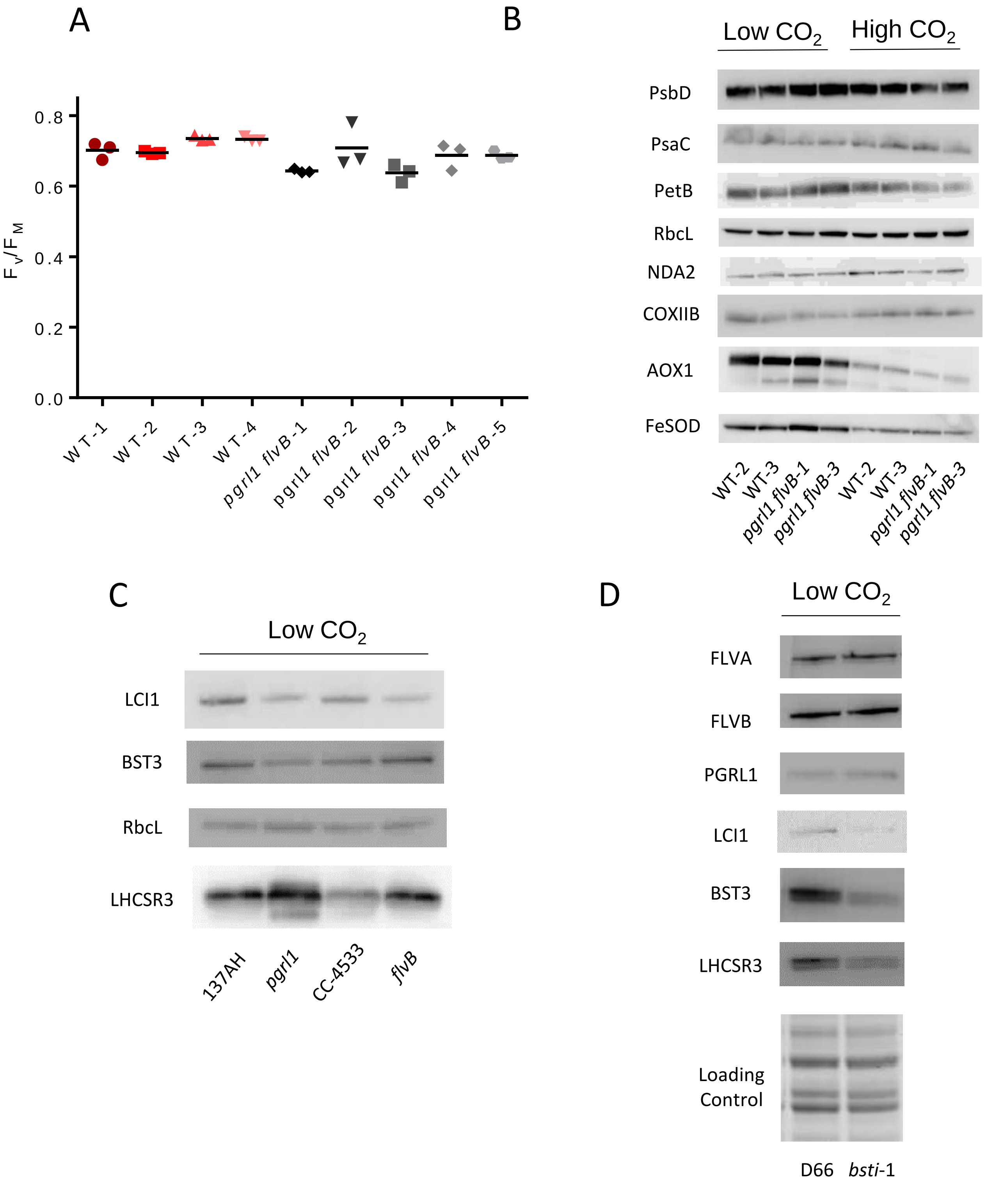


**Extended data Fig. 4. Photosynthesis and CCM components assessment.** (**A**) Maximum PSII efficiency ($F_V/F_M$) of double mutants (*pgrl1 flvB*-1 to -5) and their control strains (WT-1 to -4), measured using a PAM fluorimeter after 15 min dark adaptation. (**B**) Immunodetection of the PSII subunit PsbD, the PSI subunit PsaC, the cytochrome $b_6f$ complex subunit PetB, the large RuBisCO subunit RbcL, the type II NADH dehydroagenase NDA2, the mitochondrial cytochrome *aa3* oxidase subunit CoxIIB, of the alternative oxidase AOX1, and of the Fe superoxide dismutase FeSOD, in two independent *pgrl1 flvB* double mutants and controls grown under Low $CO_2$ or High $CO_2$. (**C**) Immunodetection of LCI1, BST3, RbcL and LHCSR3 in *pgrl1*, *flvB*, and their respective control strains (137AH and CC-4533). (**D**) Immunodetection of FLVA, FLVB, PGRL1, LCI1, BST3 and LHCSR3 in *bsti*-1 and its control strain (D66).

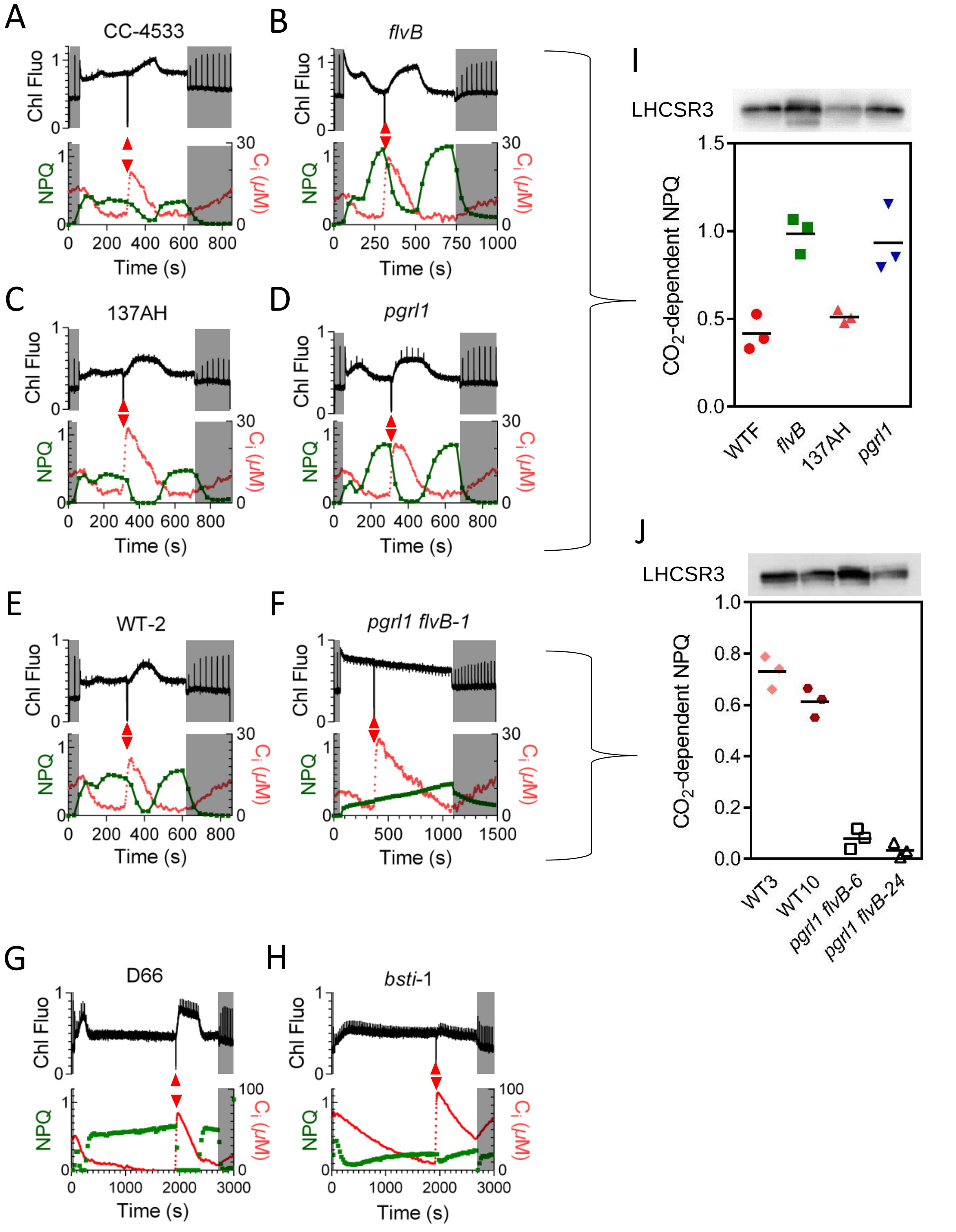


**Extended data Fig. 5. Chlorophyll fluorescence, NPQ and $C_i$ measured during dark-light-dark transients in *pgrl1*, *flvB*, *pgrl1 flvb*, *bsti-1* mutants and their respective controls.** (**A-H**) Chlorophyll fluorescence patterns (upper panels), NPQ and $C_i$ measurements (lower panels). Shown are representative experiments of three biological replicates. (**I,J**) $CO_2$-dependent NPQ was determined by subtracting the minimal NPQ obtained during the second $C_i$ depletion to the maximal NPQ obtained after the second $C_i$ depletion. Immunodetection of LHCSR3 is shown on top of each corresponding strain. Shown are mean values and replicates (n=3). All strains were grown as in **Fig. 1** under low $CO_2$.

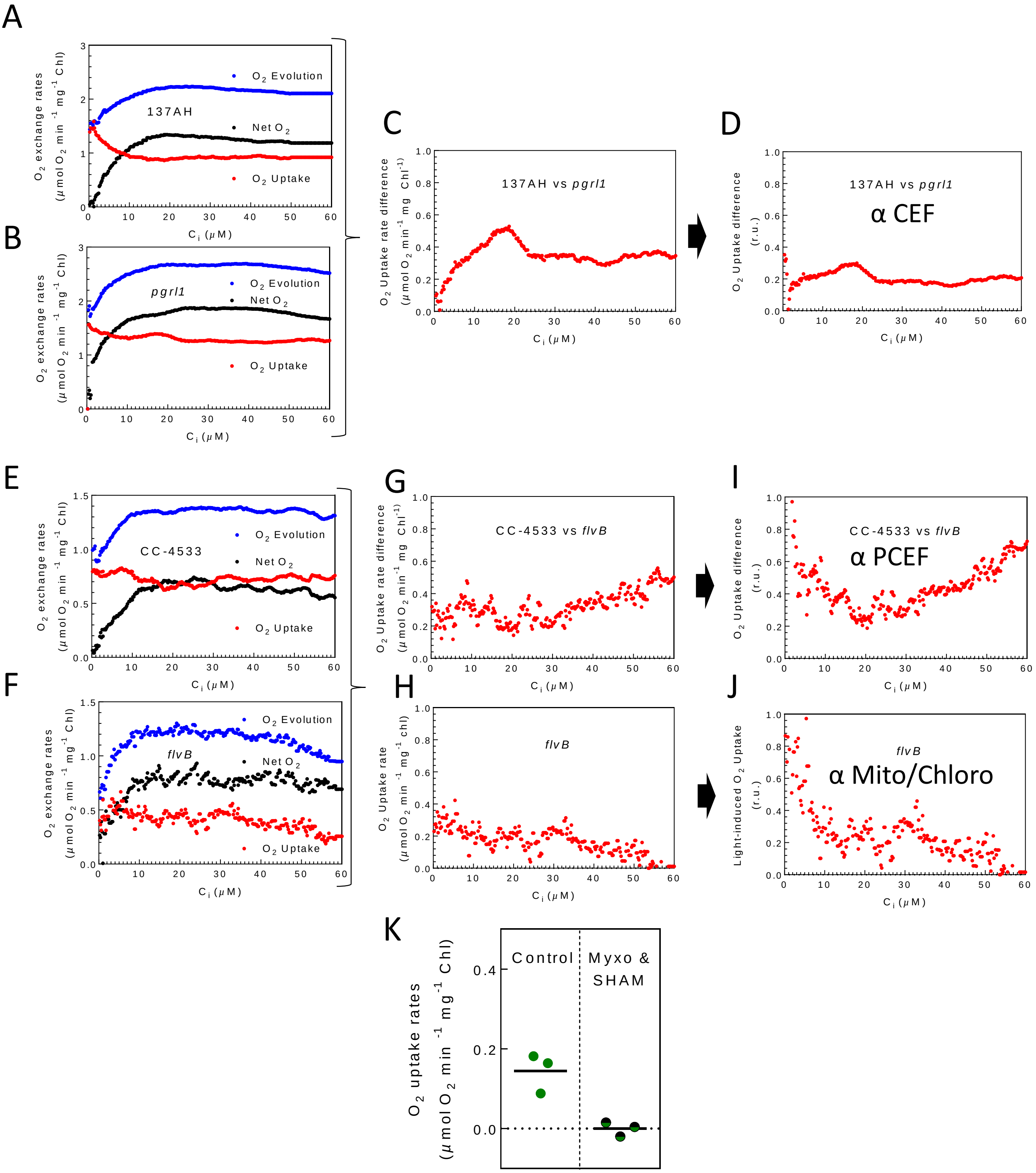


**Extended data Fig. 6. $C_i$ dependency of $O_2$ exchange rates measured by using MIMS in the presence of $^{18}O_2$-enriched $O_2$ in *pgrl1*, *flvB* and their respective controls.** Strains were grown under low $CO_2$ as in **Fig. 1**. (**A, B**) $O_2$ exchange rates in *pgrl1* and its control strain 137AH. (**C**) Difference in $O_2$ uptake rates between 137AH and *pgrl1.* (**D**) Difference in $O_2$ uptake rates as determined in (**C**) normalized to the net $O_2$ production of *pgrl1* is used to determine the CEF contribution (**E, F**) $O_2$ exchange rates in *flvB* and its control strain (CC-4533). (**G**) Difference in $O_2$ uptake rates between CC-4533 and *flvB*. (**H**) Light-induced $O_2$ uptake in *flvB* mutant. (**I**) The difference in $O_2$ uptake rates as determined in (**G**) normalized to the net $O_2$ production in *flvB* is used to determine the contribution of PCEF. (**J**) the difference in $O_2$ uptake rates as determined in (**H**) normalized to the net $O_2$ production in *flvB* is used to determine the contribution of CMEF. (**K**) Effect of the mitochondrial respiration inhibitors myxothiazol and SHAM on the light-induced $O_2$ uptake rate measured in *flvB* mutant at low $C_i$ after 3 min of illumination. Shown are mean values and replicates (n=3).

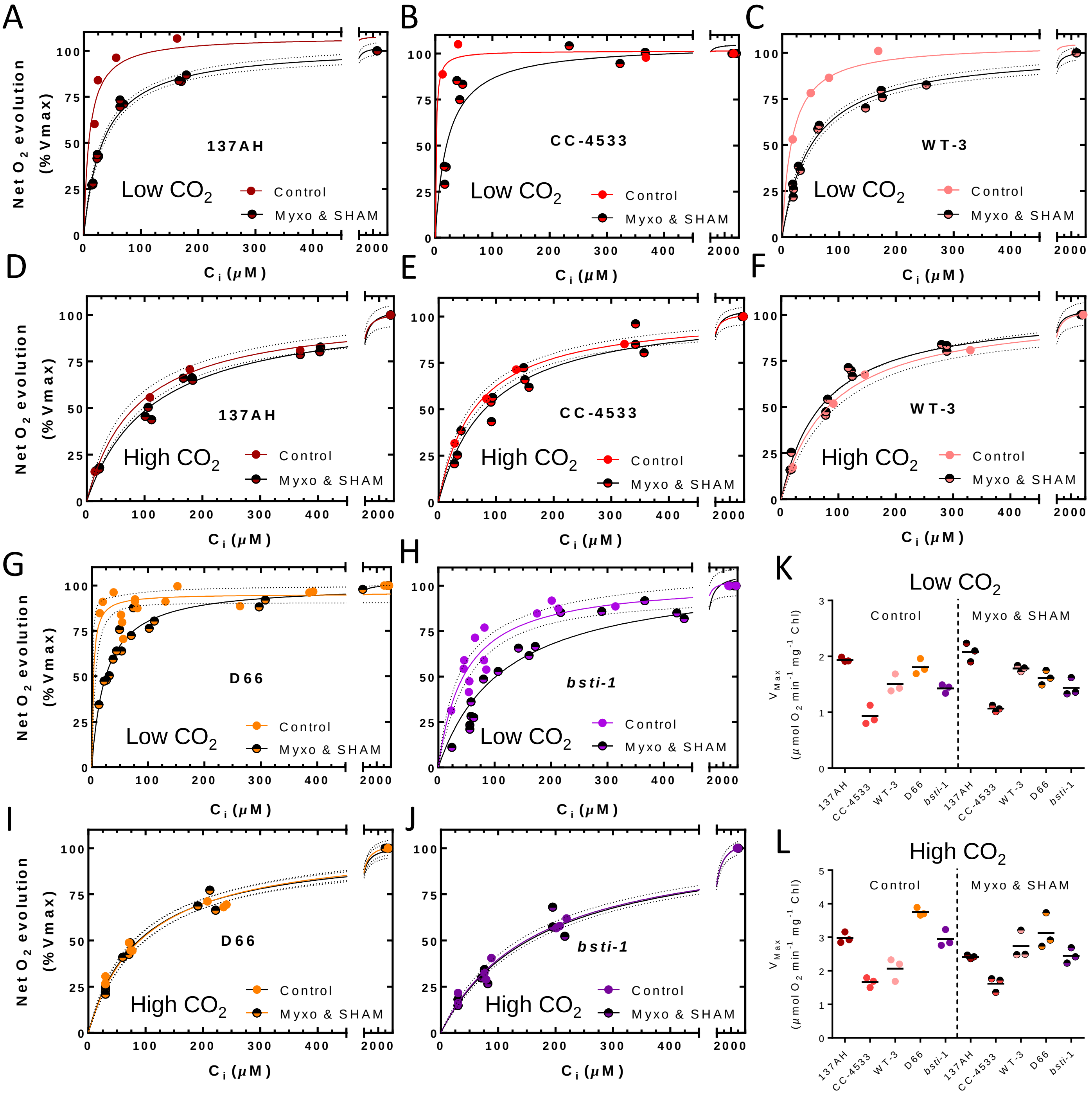


**Extended data Fig. 7. Effect of mitochondrial respiration inhibitors on the $C_i$ affinity of net $O_2$ photosynthesis.** Photosynthetic net $O_2$ production was measured as in **Fig. 1** in low $CO_2$ (**A, B, C, G, H, K**) or high $CO_2$ (**D, E, F, I, J, L**) grown cells in the absence or presence of the two mitochondrial respiration inhibitors myxothiazol (Myxo, 2.5 $\mu$M) and salicyl hydroxamic acid (SHAM, 400 $\mu$M). (**A-J**) For each replicate, net $O_2$ production was measured at four different concentrations of $C_i$ and normalized to the maximum photosynthetic net $O_2$ production. Shown are three replicates for each strain (dots) and hyperbolic fit with variability (plain lines, dotted lines) (**K, L**) Maximum net $O_2$ production rates. Shown are mean values and replicates (n=3).

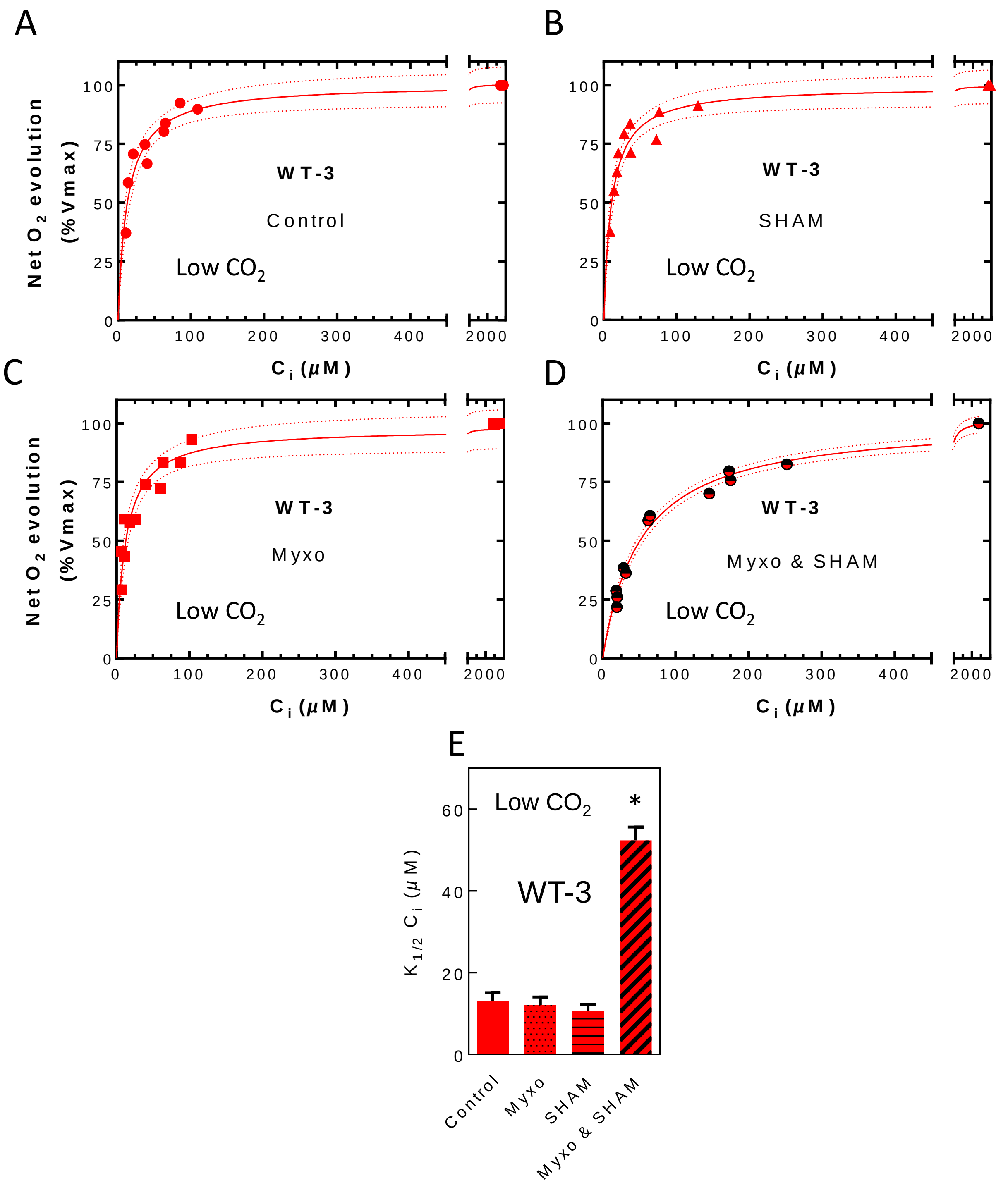


**Extended data Fig. 8. Effects of respiratory inhibitors myxothiazol and SHAM added separately or simultaneously on the $C_i$ affinity of net $O_2$ photosynthesis.** Photosynthetic net $O_2$ production was measured as in **Fig. 1** in WT-3 low $CO_2$ grown cells in the absence (**A**) or in the presence of respiratory inhibitors. (**A**) In the absence of inhibitors. (**B**) in the presence of 400 μM SHAM. (**C**) in the presence of 2.5 μM myxothiazol. (**D**) in the presence of both 2.5 μM myxothiazol and 400 μM SHAM. Shown are three replicates for each strain (dots) and hyperbolic fit with variability (plain lines, dotted lines). (**E**) $K_{1/2}$ values were determined from hyperbolic fits**.** Shown are mean values (n=3, ±SD). Asterisk represent significant difference ($p<0.05$, multiple t-test).
